# The effect of environment on the evolution and proliferation of protocells of increasing complexity

**DOI:** 10.1101/2022.07.14.499621

**Authors:** Suvam Roy, Supratim Sengupta

## Abstract

The formation, growth, division and proliferation of protocells containing RNA strands is an important step in ensuring the viability of a mixed RNA-lipid world. Experiments and computer simulations indicate that RNA encapsulated inside protocells can favour the protocell promoting its growth while protecting the system from being over-run by parasites. Recent work has also shown the rolling-circle replication mechanism can be harnessed to ensure rapid growth of RNA strands and probabilistic emergence and proliferation of protocells with functionally diverse ribozymes. Despite these advances in our understanding of a primordial RNA-lipid world, key questions remain about the ideal environment for formation of protocells and its role in regulating the proliferation of functionally complex protocells. The hot spring hypothesis suggests that mineral-rich regions near hot-springs, subject to dry-wet cycles provide an ideal environment for the origin of primitive protocells. We develop a computational model to study protocellular evolution in such environments that are distinguished by the occurrence of three distinct phases, a wet phase, followed by a gel phase, and subsequently by a dry phase. We determine the conditions under which protocells containing multiple types of ribozymes can evolve and proliferate in such regions. We find that diffusion in the gel-phase can inhibit the proliferation of complex protocells with the extent of inhibition being most significant when a small fraction of protocells is eliminated during environmental cycling. Our work clarifies how the environment can shape the evolution and proliferation of complex protocells.

## 1 Introduction

The RNA world hypothesis, according to which an RNA-based life preceded the DNA and protein-based life on prebiotic earth, has been an important hypothesis regarding the origin of life. Abiotic synthesis of ribonucleotides [1, 2, 3] in simulated prebiotic scenarios, discovery of ribozymes [4, 5, 6] and their synthesis by *in vitro* evolution processes [7, 8, 9], are circumstantial evidence that have provided indirect support for this hypothesis. The possibility of coexistence of RNA alongside amino acids [10] and lipids [11] on prebiotic earth has lead to the speculation of an updated version of the hypothesis: the RNA-lipid-peptide world. In this scenario spontaneously formed lipid vesicles [12, 13] can encapsulate RNA molecules and such encapsulation turns out to be advantageous to both RNA molecules [14, 15, 16, 17] and the vesicles that encapsulate them, as nucleotides are shown to have stabilizing effects on the lipid membranes [18]. RNA encapsulation also leads to the growth of lipid vesicles as it creates a difference in osmotic pressure between the vesicles containing RNA and empty vesicles, resulting in lipid transfer from the empty ones [19]. Presence of amino acids and small peptides is also advantageous as they, besides stabilizing the vesicle membranes [20], can turn them lipophilic [21], thereby causing further growth of the vesicles by lipid transfer from non-lipophilic vesicles. Large vesicles can divide when subject to external forces [22, 23] and distribute their contents into daughter vesicles.

Computer simulations have shown that the evolution of functional RNA sequences within protocells offers several advantages [24, 25, 26, 27, 28] over their evolution in spatially open systems [29, 30]. Ma and collaborators [26] have used Monte Carlo simulations to show how ribozymes created in open spatial systems can be engulfed by protocells, conferring a selective advantage to ribozyme containing protocells that allows such protocells to survive and increase in number even as the ribozymes created in the open spatial system are gradually eliminated. Higgs and collaborators [27] have demonstrated that protocells can tolerate a lower replication rate as well as lower replication fidelity of replicases without being overrun by selfish genetic elements. Even though these computational models provide important insights into the dynamics and evolution of RNA strands and protocells in a primordial RNA world, a plausible mechanism of formation of long and potentially functional RNA strands from basic building blocks is not discussed. These issues were addressed in [31] where we showed that both concatenation and template-directed primer extension in an environment subject to dry-wet cycling is essential for creation of long and structurally complex RNA sequences from activated nucleotides. Our work raised the question of what might happen if such processes are confined within vesicular membranes. We have recently shown, using realistic, experimentally determined parameters, that a population of small vesicles encapsulating a small number of RNA molecules initially can evolve to a population consisting of large protocells containing multiple types of ribozymes via non-enzymatic rolling circle replication mechanism, protocell division and preferential selection of vesicles with higher number of RNA strands inside them [32]. Thus, a mixed RNA-lipid-peptide world can eventually lead to the possible emergence of primitive protocellular life with protocells gradually increasing in complexity through the stochastic creation of a diverse set of ribozymes [32, 33]. A key element, omitted in the above discussion, is that of the plausible environment for the origin and spread of primitive protocells. There has been much speculation on the ideal environmental conditions since it continues to be a critical factor in determining the viability of any protocell model of origin and evolution of life.

There are two major hypotheses regarding a suitable environment for life’s origins: the hydrothermal vent hypothesis [34] and the hot spring hypothesis [35]. There have been significant effort in analysing the efficacy of both these scenarios. The temperature gradient of submarine hydrothermal vents have been shown to be useful for RNA polymerization reactions in creating long RNA molecules [36, 37, 38, 39]. Normally in aqueous solutions the average length of polymers depend on the polymerization (*K*_*pol*_) and hydrolysis rates (*K*_*hyd*_) as 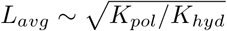 [40]. But in presence of a temperature gradient molecules drift along the temperature gradient, a phenomenon known as thermophoresis [41]. It causes an influx of monomers and as a result the average length in this case also depends on the monomer influx rate (J) as 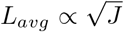 [36]. Therefore the temperature gradient of submarine hydrothermal vents can generate much longer RNA polymers compared to normal aqueous solutions [37]. Nevertheless, the hydrothermal vent hypothesis has certain disadvantages. The higher concentration of ionic solutes in seawater inhibits the formation of lipid vesicles and encapsulation of polymers inside them [42]. Therefore submarine hydrothermal vents, even though suitable for formation of long polymers, are not the ideal environment for formation of a protocellular life. Proponents of the alternative hot spring hypothesis argue that the region around hot springs/geothermal pools provides the ideal environment for the origin of life and evolution of protocells [35, 43]. Besides being rich in elements necessary for prebiotic chemistry [44, 45], the periodic dry-wet cycles in such regions help in creation of longer polymers [46, 31]. Lipid molecules spontaneously formed in such regions can assemble into vesicles [47]. During the dry phase, vesicles fuse into multilamellar structures and RNA monomers and oligomers get trapped between different layers of the lamella. The reduced water activity in this state aids phosphodiester bond formation and thereby helps in synthesis of long RNA polymers at rates faster than the hydrolysis and degradation rates that break up such polymers. On the advent of the subsequent wet phase the lamella swells into vesicles while still containing the RNA polymers and as a result the polymers get encapsulated in the vesicles [35, 48]. An intermediate phase between the wet and dry phase results when the vesicle membranes start to fuse, thereby creating channels between them, allowing for free long-range movement of both monomers and long polymers that were previously confined to a vesicle [35].

Along with laboratory-based experiments, recently few experiments done near hot springs have provided confirmation of the spontaneous assembly of vesicles and formation of long RNA polymers in such locations [42, 49, 50]. All of these experimental findings, while providing strong empirical support of plausible prebiotic processes are not adequate in explaining how primitive protocellular life could have evolved and spread in such regions. The experimental investigations in real hot spring environments are still at a stage of infancy and it is indeed extremely difficult to conduct experiments of this scale in such inhospitable environments. In this context, computational models can be very useful in testing many of the key conceptual issues on protocell evolution in such environments that have been the subject of considerable speculation in the literature.

In our previous work [32] we considered the rolling circle replication process of circular RNA strands inside lipid vesicles. That choice was based on its effectiveness in maintaining exponential growth of the number of RNA strands over a large temperature range [32]. If the hot spring hypothesis is true a similar mechanism should be applicable even in such environments. Environmental factors like wet-dry cycling and its impact on the physiology of the lipid lamella were not considered in our previous study. Here we specifically test this key feature of the hot spring hypothesis of the origin of life by studying how phase transitions from the wet to gel-like and subsequently to a dry phase change the mobility of large RNA strands and consequently affect the evolution of the protocell population. While placing our model in a hot-spring environment we segregated the rolling circle replication and vesicle division processes in the dry and wet phases respectively. This is because lack of water inside the lamellae results in a kinetic trap [51], concentrating reactants and enhancing the rates of different types of polymerization reactions. In contrast, the wet phase causes vesicles to bud off from the lamella, increases their mobility, which in turn subjects them to external stress and shear forces that can lead to division of larger vesicles. We considered a 2D lattice to represent the lamella and assumed spontaneous formation of vesicles from different lattice sites during the wet phase, with vesicles formed at each site encapsulating the RNA molecules present at that site. In light of the increased mobility of long polymers during the gel phase, we included an extra process of long polymer diffusion from one site to another during this phase when the vesicles start to fuse.

We found that in the presence of dry and wet phase only, the initial population of protocells can evolve towards a state where protocells containing multiple types of ribozymes dominate. However, when the process of long polymer diffusion in the gel phase is taken into account, there is a significant slow down in the evolution of the protocell population, suggesting an inhibitory effect of gel phase diffusion. This effect is most pronounced when we consider degradation of entire protocells in the wet phase with small probabilities. The long polymer diffusion process in the gel phase then becomes the primary reason for gradual decay and ultimately elimination of RNA strands and as a result the protocell population fails to evolve. On the other hand, dry and wet phase alone or a dry, wet and gel phase that does not facilitate long polymer diffusion, can sustain the evolution even in presence of protocell degradation process. Finally in presence of dry and wet phase and absence of diffusion in the gel phase we also observed spatial expansion of the protocell population when initially one or very few of the sites contain circular templates, even when protocells are allowed to degrade in the wet phase. The growth of RNA strands inside those protocells followed by their subsequent division and occupation of neighboring sites by daughter protocells eventually lead to the outward spread of the protocell population with multiple ribozymes until it covers almost the entire lattice. Our work provides quantitative support for the viability of the hot spring environment in creating and sustaining an evolving population of protocells characterized by increasingly complex functionality.

## 2 Methods

According to the hot-spring hypothesis [43, 35] lipid molecules can periodically organize themselves to form three different types of structures determined by environmental conditions. In the dry phase they can form a multilamellar structure on mineral surfaces. Such structures act like kinetic traps confining reactant molecules and promoting polymerization reactions with activated nucleotides [31, 52]. In the wet phase, local regions of that lamellae can swell into vesicles that can confine long RNA sequence fragments and in the gel phase the vesicle membranes fuse creating connected channels that can allow for unrestricted movement of long polymers across the entire region. To simulate such a scenario, we start with a 2D square lattice (of size *N* × *N* = 30 × 30) on a mineral surface representing the lamella. Each square lattice site (of size Δ*x* ~143 nm) provides a site for polymerization reactions to occur while boundaries between lattice sites prevent free movement of long polymers in the dry phase. In the wet phase, the lamellae from each site can bud off into different vesicles thereby providing a confining environment that traps RNA polymers within the protocells. This is implemented by restoring the boundaries associated with each lattice site thereby ensuring that each site acts as an independent protocell during the wet phase. In the gel-phase, the disappearance of the boundaries between the lattice sites is mimicked by allowing for free movement of long polymers across the entire lattice. We first start with one random circular ssRNA (denoted by *s*) template of length 200 nt [53] at each lattice site, at the beginning of a dry (lamellae) phase. Subsequently, we also consider an initial scenario where a single site has one or a small number of circular templates while the remaining sites are devoid of any such templates. During the dry phase the molecules have very low mobility with only monomers and small oligomers (≤ 8 nucleotides) capable of diffusing freely across the lattice. The longer circular strands will remain fixed at their respective lattice sites. At each lattice site inside the lamellae, a circular ssRNA can transform into a circular double stranded RNA (dsRNA denoted by d in equations below) by attaching with a small complementary primer (~ 8 nt) followed by template-directed extension of the primer. Upon becoming full length the primer can extend further along the template by displacing its other end from its initial point on the template, thereby creating a hanging chain. When the hanging chain attains a length equal to the template, it cleaves and gets separated from the circular dsRNA [32]. This process, called the rolling circle replication process [54, 55, 56], have been shown to be more effective [28] in producing long ssRNA molecules and avoid the strand-separation problem associated with double-stranded RNA sequences. The rate of creation of circular dsRNA from circular ssRNA and the rate of rolling circle replication of circular dsRNA is taken to be random for random templates and sampled from the distribution [32] *K_rep_* = *e*^−3.22–0.005×*L*–2.8×*rand.norm*(0.35,0.0667)^ *h*^−1^ (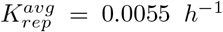 and 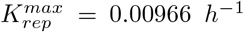 for *L* = 200) that was determined using the experimental primer extension rates [52, 57, 31]). The newly created, open-ended single strands (denoted by l) are diverse in terms of secondary structures [32] because of the error-prone nature of the replication process[52, 57]. Therefore we assume a small fraction of them will attain catalytic capabilities associated with different types of ribozymes like the replicase (r), cyclase (c), nucleotide synthase (n) and peptidyl transferase (p). A replicase can catalyze the process of circular ssRNA to circular dsRNA formation by the template-directed primer extension mechanism and subsequently from circular dsRNA to an open-ended ssRNA by the rolling circle mechanism. At a site ‘ij’ (i.e. the site labelled by the row-index ‘i’ and column-index ‘j’; 1≤ *i ≤ N*, 1 ≤ *j ≤ N*) the rate of these processes would be *K*_*fast*_*s*_*ij*_*r*_*ij*_*/V* and *K*_*fast*_*d*_*ij*_*r*_*ij*_*/V* (*K*_*fast*_ = 0.362 *h*^*−*1^ [32]), where we divide by a volume factor *V* = 100 for each lattice site in unit of strand numbers to match the dimension of these 2nd order reactions. A cyclase (a type of ligase) can ligate the open ends of a non-catalytic open-ended ssRNA creating a circular ssRNA at a rate *K*_*fast*_*c*_*ij*_*l*_*ij*_*/V*. A nucleotide synthase can create new free nucleotides in a monomer deficient system. Therefore in the monomer limited scenario that allows for the possibility of creation of a nucleotide synthase, we multiply each replication rate of site ‘ij’ with a term 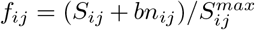, where 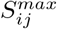 and *S*_*ij*_ are the maximum and instantaneous number of monomers in a site in units of 200-mers and b is the number of monomers in units of 200-mers a single nucleotide synthase can create (we set *b* = 1 and 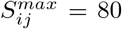 in the simulations). Finally, a peptidyl transferase can join free amino acids that are assumed to be present in the system, to create small peptide chains. During the wet phase when the lamellae from different lattice sites swell into vesicles, such small peptide chains can bind with the lipid membrane of the vesicles and turn them into lipid attracting membranes, thereby causing them to grow in size [21] increasing the threshold volume (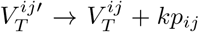, where *k* is the strength of a peptidyl transferase, taken to be 20) beyond which the vesicle divides into two daughter protocells. Finally all types RNA molecules inside the lamellae can degrade at a certain rate (h). These reactions can be represented as a set of equations:

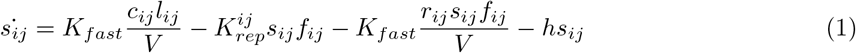

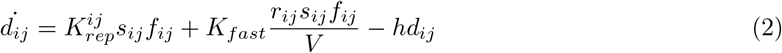

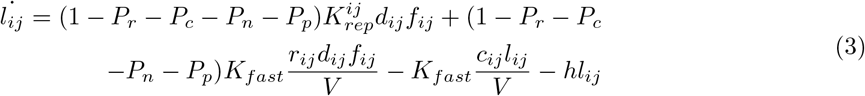

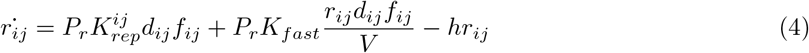

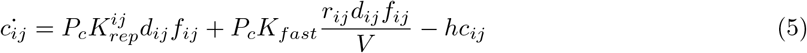

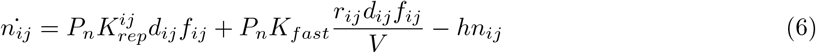

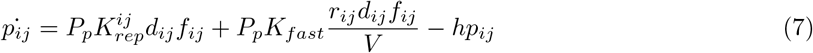

We solve the fully stochastic version of these 7 equations [32] in our simulations (more details are given in the supplementary information file.). As a consequence of these above mentioned reactions (which are favored by the low amount of water in the dry phase), the number of strands at each lattice site can grow and eventually be encapsulated in a vesicle forming at that site from the lamella during the subsequent wet phase.

Presence of high amount of water in the wet phase will cause the vesicles to undergo Brownian motion, thereby making them highly mobile. For example a vesicle of diameter 60 nm can move across our entire lattice in less than 10 sec [58]. Such high mobility of the vesicles can make them collide with sharp rock edges or move through rock pores. They can even be subjected to shear and stress due to turbulent flow of water in which they are immersed. Such forces can induce division of larger vesicles [22, 23]. In our model, when the number of RNA molecules inside such a vesicle exceed a certain threshold (depending on the number of peptidyl transferase ribozymes inside it), the vesicle will divide into 2 daughter vesicles. As division causes an increase in the total surface area (26% for perfectly spherical vesicles), we assume the elimination of another vesicle in the process to provide the lipid molecules needed for that extra surface area. Selection is introduced by imposing the condition that vesicles with fewer RNA strands inside them are more likely to get eliminated. Therefore we take the elimination probability of a vesicle to be proportional to the difference between the number of RNA strands inside it and in the dividing vesicle.

In the dry phase, there are no vesicles and long RNA strands on specific site are immobile even though monomers and short oligomers can diffuse freely. During the transition from the wet to the dry phase, as the excess water starts to dry up due to increasing temperatures, creating a gel phase, it causes the vesicle membranes to fuse, creating channels from one vesicle compartment to another on the multi-lamellar lattice. This phase allows for increased mobility of even large RNA strands, thereby allowing for their potential dispersal to different regions of the lattice. The diffusion coefficients of the long strands can be related to their hopping probabilities from one lattice site to a neighboring site by the equation [59]:

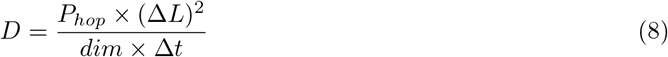

Diffusion of molecules from one site to another also depends on the difference in the number of strands between them. The molecules from the central site are more likely to diffuse to the neighbor that is relatively emptier. The size of the lattice site is Δ*L* = 143 *nm*. Therefore for a time step size Δ*t* = 0.008 *hr* the maximum diffusion coefficient (for *P*_*hop*_ = 1) in 2D (dim=2) would be *D* ~ 1.28 *μm*^2^*h*^*−*1^, which is ~ 10^5^ times smaller than the diffusion coefficients of 200 mer RNA strands (single and double stranded) if they were present freely outside the vesicles in the fully hydrated wet phase [60, 61]. Although experimentally it is found that double strands have lower diffusion coefficient than single strands, we found that our results do not differ even if we consider different hopping probabilities for the circular dsRNA molecules. We take the duration of the dry, wet and gel phase to be equal to 8 hours each, so that they can appear periodically for each day of length 24 hours. The simulations were run for many such days using periodic boundary conditions on the lattice.

## 3 Results

### 3.1 Effect of the individual phases

We first simulated the evolution of RNA strands and protocells in the dry and wet phase after inactivating the gel phase in order to understand the dynamics in the absence of long-strand diffusion. Such a scenario also provides a benchmark for evaluating the impact of the gel phase when it is subsequently switched on. We find that the number of strands in a few sites which happen to contain faster replicating circular templates initially, grows and get encapsulated in vesicles during the subsequent wet phase. However, at most other sites the initial circular templates have low-medium replication rates as a result of which new RNA strands including ribozymes are not produced at a rate that is faster compared to the degradation process. As a consequence, those sites become devoid of RNA strands. This results in the concentration of a large number of strands at a few sites while most of the remaining sites become empty. Eventually, as the vesicles originating from those few sites start to divide when the threshold volume is attained, the newly created daughter vesicles occupy the empty nearest neighbor sites. Through this vesicle division process, the RNA molecules gradually spread over the entire lattice and we find that ~ 90% sites (and subsequent protocells) contain all 4 types of ribozymes at equilibrium (Fig-2, D=0). The evolutionary push towards dominance of protocells with all 4 types of ribozymes comes from the formation of a cooperative network between these 4 types of ribozymes [32]. The supplementary video Vid S1 shows the evolution of different types of RNA strands and ribozyme diversity across the entire lattice (more details about the video are given in the Supplementary information).

**Figure 1:**
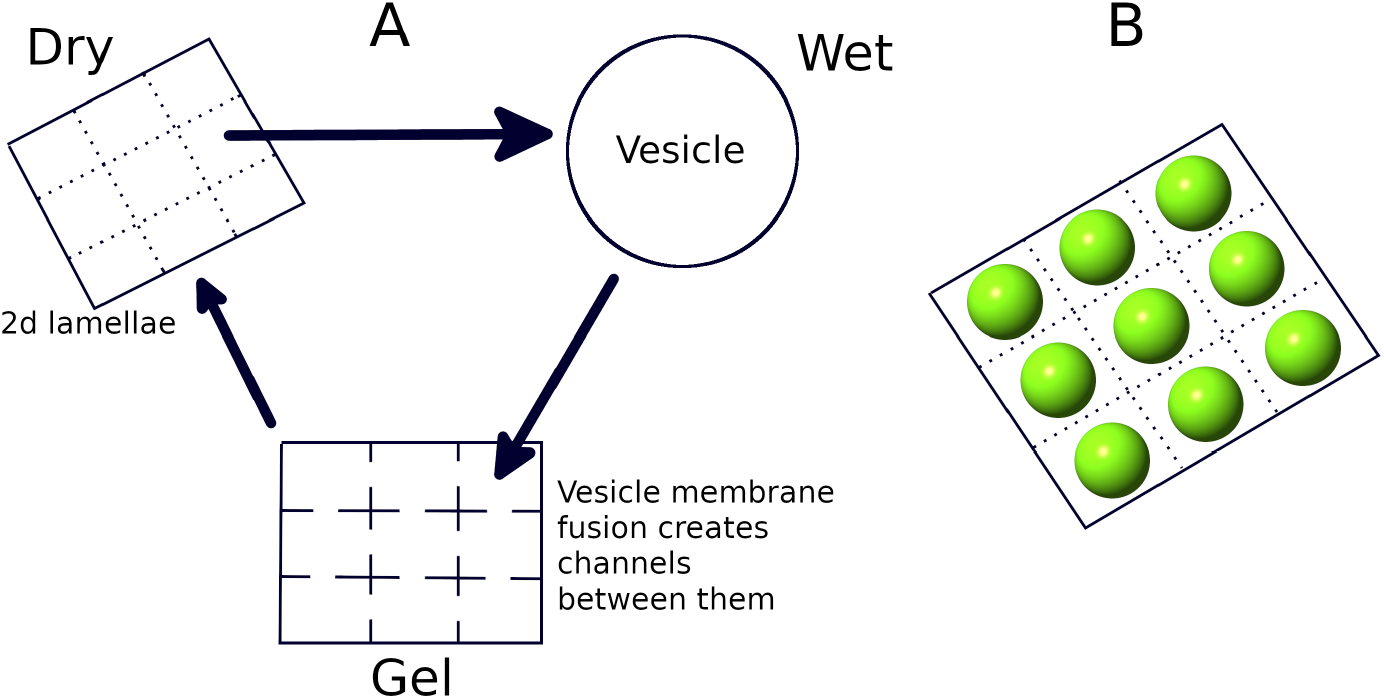
**A:** Pictorial representation of the 3 phases that periodically occur in hot spring environments. Multilamellar structures can form on mineral surfaces from lipid molecules in the dry phase, one layer of which is represented as a 2D lattice containing sites for RNA polymerization. Each site can swell into a vesicle in the wet phase. In the gel phase the vesicles get deposited on the 2D surface and their membranes start to fuse, creating channels between them that can allow for long-range diffusion of large RNA strands. Subsequent to this stage, the multilamellar structure forms again in the next dry phase. **B:** 3D representation of formation of vesicles from the lamella in the wet phase.

**Figure 2:**
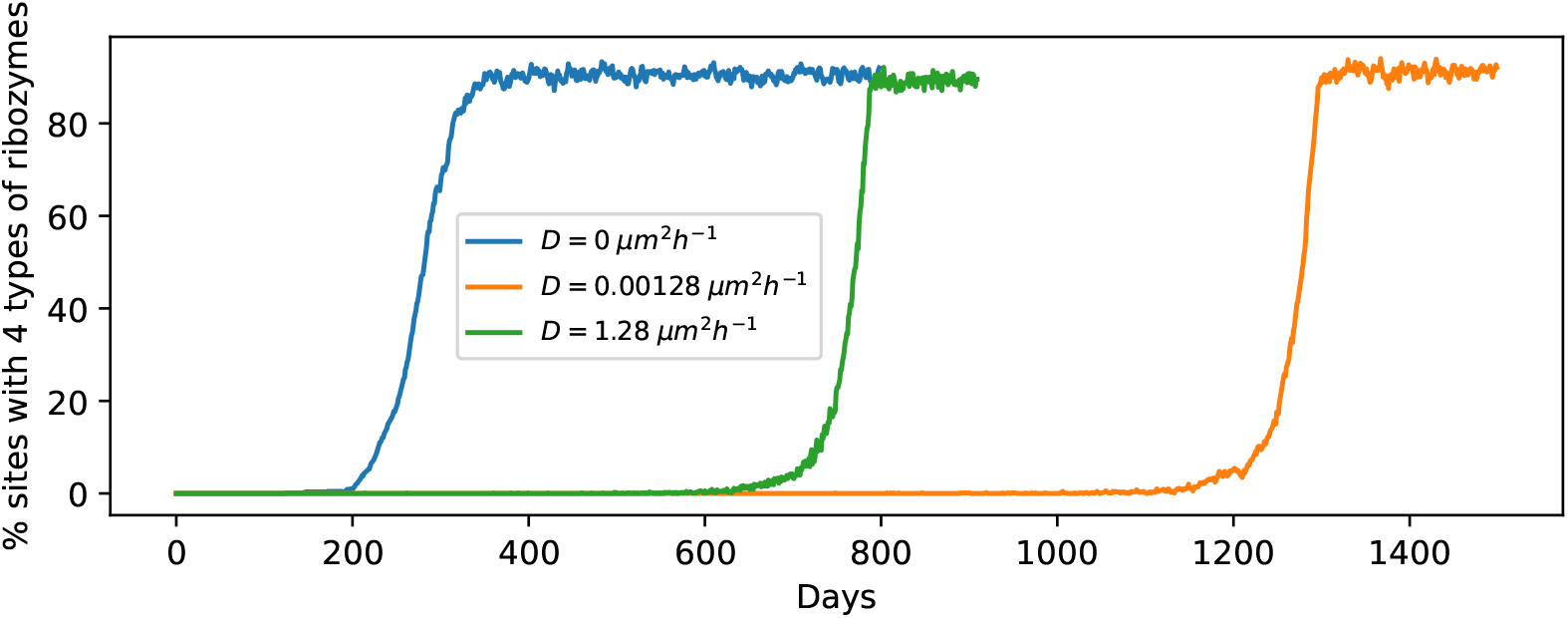
Time evolution plots (1 trial each) showing the percentage of sites containing all 4 types of ribozymes for *D* = 0 *μm*^2^*h*^*−*1^ (blue); *D* = 0.00128 *μm*^2^*h*^*−*1^ (orange) and *D* = 1.28 *μm*^2^*h*^*−*1^ (green) starting from 1 circular ssRNA per site initially.

When the gel-phase is switched on, allowing for diffusion of long RNA polymers in that phase, the evolution towards equilibrium is much slower. The plots for the percentage of sites with 4 types of ribozymes vs time for 2 diffusion coefficients are shown in Fig-2. Unlike the D=0 case, the time at which the curve starts rising steeply from 0 occurs at different times for different trial runs even for the same diffusion coefficient. Nevertheless, we observed that the time evolution for non-zero D values are still much slower than the D=0 case across all trials. We therefore show the curves for only 1 trial for each diffusion coefficient in Fig-2. The video Vid S2 shows the proliferation of RNA strands across the entire lattice when diffusion is allowed in the gel phase (for *D* = 1.28 *μm*^2^*h*^*−*1^). The dispersal of long RNA strands can impede the growth of RNA strands at a specific site, delaying the protocell division process and eventual proliferation of daughter protocells to neighbouring sites. For example if a replicase emerges at a site having a faster replicating template initially, then it would speed up the replication process at this site by many folds thereby increasing the likelihood of formation of new ribozymes. However, if such a replicase responsible for rapid growth of RNA strands at a site were to disperse to another site in the gel phase, it would limit the subsequent growth of RNA strands and stochastic creation of ribozymes at the original site. Moreover, the effectiveness of the replicase at the new site might be hindered by the presence of fewer circular templates and/or other types of ribozymes. Similarly, if a cyclase, responsible for catalysing the formation of new circular templates on which a replicase can act to speed up the replication process, diffuses to another site, it can significantly slow down the creation of new strands at the original site. Even though long RNA strand diffusion in the gel phase has an inhibitory effect on the evolution and proliferation of protocells with increasing complexity, given sufficient time, complex protocells containing multiple ribozymes can still proliferate and occupy ~ 90% sites as evident from the equilibrated values shown by the orange and green curves in Fig-2.

### 3.2 Relocation of protocells in the wet phase

As mentioned in the methods section, the presence of water in the wet phase will cause the vesicles to undergo rapid Brownian motion. Therefore a vesicle which originates from a site on the lattice, will not deposit on the same site after the wet phase. Brownian motion of vesicles during each wet phase will effectively result in random relocation of the vesicles on the lattice. To account for this phenomenon, we randomly shuffle the vesicle positions on the lattice at the beginning of each wet phase. Vid S3 shows the evolution of protocell population in this scenario. When such relocation of protocells is allowed, the percentage of sites with four types of ribozymes takes somewhat longer to saturate (figure not shown), but not as long as it takes to reach equilibrium in presence of gel-phase strand diffusion. However, the equilibrium abundance of protocells with four types of ribozymes drops to ~ 72%. This happens because in the *absence* of relocation few initial vesicles with faster replicating strands cross the threshold volume leading to vesicle division which spreads some of their contents encapsulated in newly formed daughter protocells to neighboring sites. This process happens from a few locations on the lattice and as the process continues, it leads to the formation and outward growth of clusters of vesicles containing all four ribozymes. This happens because dividing vesicles located on the edge of the cluster are more likely to contain larger numbers of ribozymes with one of the daughter protocells more likely to eliminate vesicles neighbouring the boundary of the cluster (that have relatively fewer number of ribozymes) than those in the interior or edge of the cluster (that have relatively larger number of ribozymes). Eventually, spread of functionally complex protocells occurs through merging of these expanding clusters. This is evident from Vid S4 which tracks the evolution of protocells containing three and four ribozymes; as well as dividing protocells (see supplementary information for details). However, when protocells are relocated in the wet phase, cluster formation of protocells containing larger number of ribozymes is inhibited and the spread of complex protocells occurs due to the creation of such protocells, initially sparsely but eventually more uniformly, across the spatial landscape. This can be seen in Vid_S5 which reveals the contrasting evolutionary dynamics in comparison to the one depicted in Vid_S4.

### 3.3 Degradation of entire protocells in the wet phase

So far we have assumed that the protocells remain intact during the wet phase but in reality some protocells will degrade in every wet phase. To account for this effect, we include the process of protocell destruction in our simulation at the beginning of every wet phase. This is likely to inhibit the spread of protocells of increasing complexity in a manner that is dependent on the protocell killing probability. Each protocell can degrade along with its contents with a probability *P*_*kill*_ with the location of the degraded protocell remaining empty until it is occupied by another protocell as a result of diffusion or a division event at a neighboring site. We first consider the case without strand diffusion in the gel phase, with our simulations being initiated with 1 circular template per site. We varied *P*_*kill*_ and found that the protocell population can evolve up to a threshold *P*_*kill*_ ≤ 0.01, above which all RNA strands die out across the lattice. This threshold increases to *P*_*kill*_ ≤ 0.02 when we start with 5 random templates per site. However when the strand diffusion process is present, even for diffusion coefficients as low as 3.2 × 10^*−*5^ *μm*^2^*h*^*−*1^ (with *P*_*kill*_ = 0.01), the protocells containing RNA strands (even excluding ribozymes) can no longer be sustained in the population. The videos showing the temporal evolution for *P*_*kill*_ = 0.01 in the absence and presence of diffusion in the gel phase (corresponding to the orange and red curves in Fig-3) are provided in supplementary information (see Vid S6 and Vid S7 respectively). We arrived at a similar result even when we started the simulations with 5 templates per site. Therefore, consistent with the results of previous sections, the strand diffusion process in the gel phase is found to be counter-productive for the proliferation of protocells containing functionally complex components. Fig-3 shows the plots for the fraction of empty sites with time for three different *P*_*kill*_ values in the absence and presence of diffusion in the gel phase. We also checked the time evolution for diffusion coefficients that are 10-fold higher (*D* = 12.8 *μm*^2^*h*^*−*1^) and for experimental diffusion coefficients in hydrated conditions, which corresponds to well mixed limit in our setup. Even for those scenarios, the behaviour is similar to that observed for *D* = 1.28 *μm*^2^*h*^*−*1^ in Fig-3. Protocell relocation in the wet phase however does not hinder the evolution as much as gel phase strand diffusion does, because the protocells with all 4 types of ribozymes still dominate with an abundance of ~ 65% even when the protocells are allowed to degrade with probability *P*_*kill*_ = 0.01.

**Figure 3:**
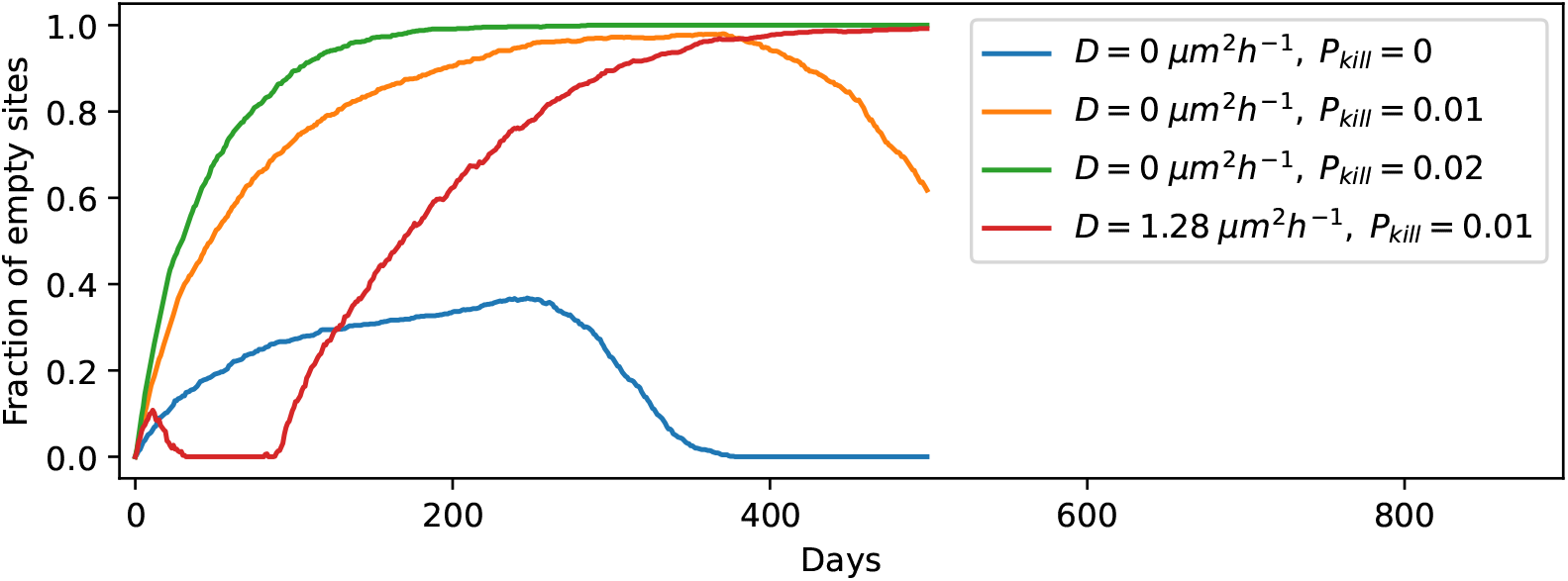
Time evolution of the fraction of empty sites for **A:** *D* = 0 *μm*^2^*h*^*−*1^ with no protocell degradation in the wet phase; **B:** *D* = 0 *μm*^2^*h*^*−*1^ when protocells degrade with probability *P*_*kill*_ = 0.01; **C:** *D* = 0 *μm*^2^*h*^*−*1^ when protocells degrade with probability *P*_*kill*_ = 0.02; **D:** *D* = 1.28 *μm*^2^*h*^*−*1^ when protocells degrade with probability *P*_*kill*_ = 0.01.

We also plot the total number of strands vs time of a random site and its 8 nearest neighbor for the case of D=0 and *P*_*kill*_ = 0.01 (Fig-4). As evident from the figure, during this time period, the total number of strands of a site goes to zero six times (counting across all the 9 sites) as a result of the protocell degradation process. The large fluctuations in the number of RNA strands observed in the plots is a consequence of the protocell division process when a parent protocell divides on reaching the threshold volume and its strands strands are divided roughly equally between its 2 daughter protocells. For example, the central site which does not contain any strands initially, acquires strands by protocell division process from either the protocell at the bottom center panel or the protocell at the bottom right panel, both of which divide in the previous wet phase. Tracking the temporal variation in a specific ribozyme (such as the cyclase) across these 9 sites reveals features that are not evident from Fig-4. Not surprisingly, the number of cyclase goes to zero more often than the total number of strands. However, this often happens as a consequence of the asymmetric distribution of RNA strands across the two daughter protocells following a protocell division event. To highlight this fact, Fig-5 shows the temporal variation in the number of cyclase for 3 of the 9 sites depicted in Fig-4. In Fig-5(B) (central site) when the cyclase number goes to 0 from 6 just after *t* = 450, the number of cyclase at a neighboring site (Fig-5(C)) becomes 6, indicating that one of the daughter protocells of the central site that acquired all the cyclases after division has occupied a neighboring site while the other daughter cell which does not contain any cyclase has replaced the parent protocell at the central site.

**Figure 4:**
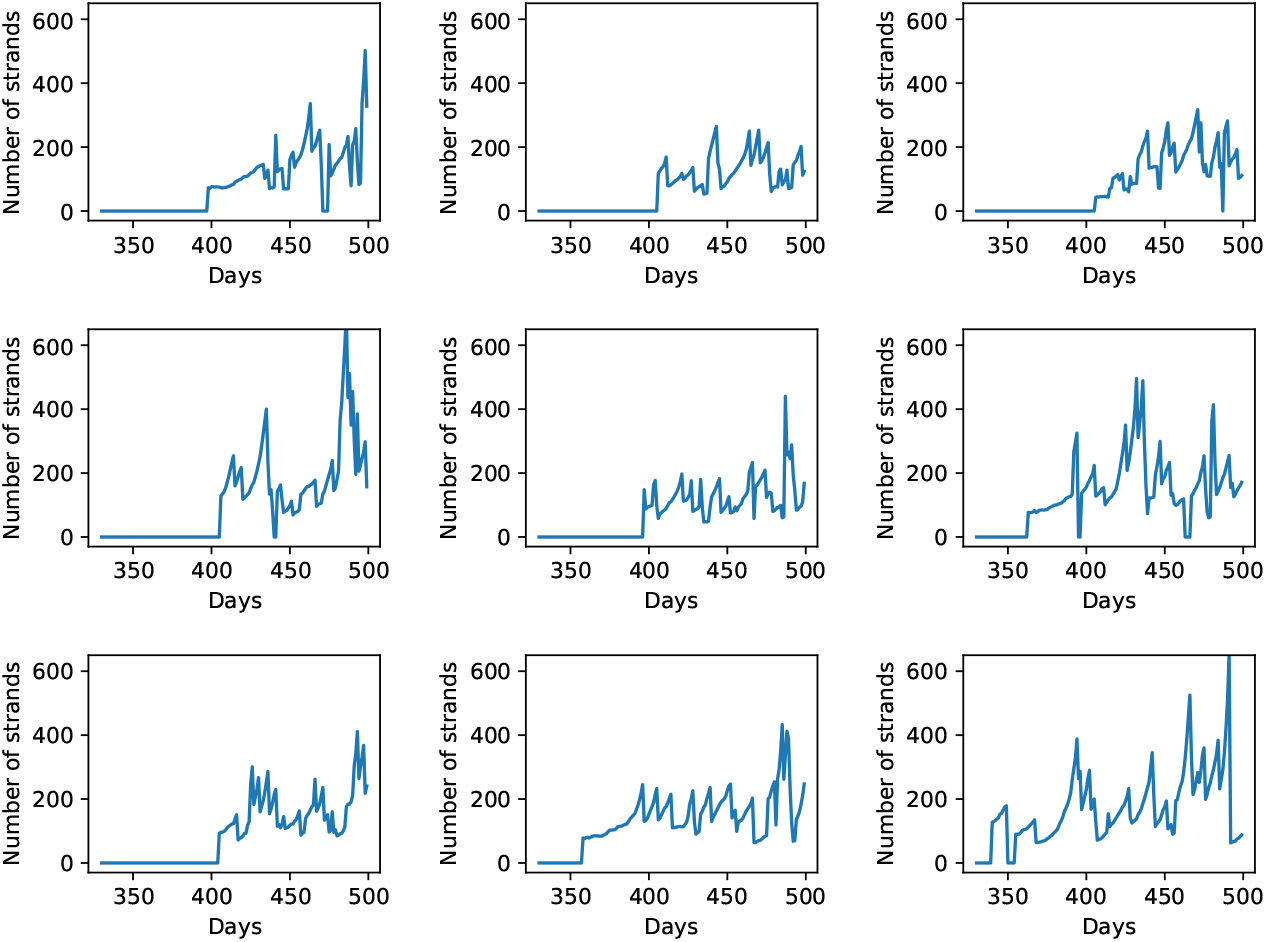
Total number of RNA strands vs time for a site having a low *K*_*rep*_ value of the initial template (middle row, central panel) and its 8 neighboring sites for the case when protocells can degrade in the wet phase with probability *P*_*kill*_ = 0.01.

**Figure 5:**
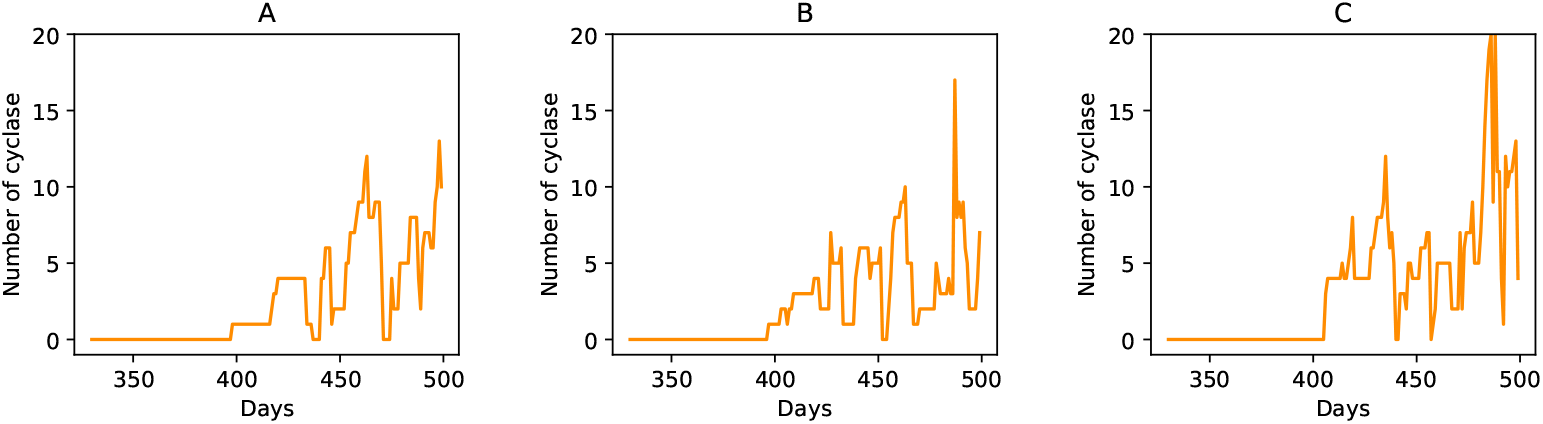
Number of cyclases vs time for sites corresponding to **A:** top-left site, **B:** middle-central site and **C:** the middle-left site; shown in Fig-4

### 3.4 Spatial expansion of protocell population

Given how a protocell can divide and its daughters can occupy a nearest neighbor site, we wanted to ascertain if the protocell population can proliferate across the entire lattice, starting from an initial state where only a few sites contain the initial templates. We first checked for the case with strand diffusion in the gel phase (*D* = 0 *μm*^2^*h*^*−*1^), protocell relocation and degradation processes were all absent. We varied the fraction of initial sites that contain a circular ssRNA and found that the protocell population can evolve and spread to the entire lattice if at least 2% of initial sites contain a template. Interestingly even if we start with only 1 non-empty site (out of 900 sites) the protocell population can still evolve and spread, provided that the single site contains at least 5 circular templates with *K*_*rep*_ values taken from the distribution. Fig-6 shows the population evolution for this case at 3 different stages. Next we included the strand diffusion process (with *D* = 0.00128 *μm*^2^*h*^*−*1^) in the gel phase. We found that at least 20% sites need to contain at least 5 templates for the protocell population to evolve and expand spatially. Otherwise the entire RNA population gradually dies out. In the case of only *one* non-empty site, even starting with 50 templates is inadequate to ensure the proliferation of protocells when strand diffusion is present. Therefore, strand diffusion in the gel phase can prevent the gradual spatial expansion of functionally complex protocellular population. However when the protocell relocation process in the wet phase was allowed to occur instead (with strand diffusion turned off), 2% initial sites containing 1 template each were able to evolve and expand, indicating this process does not hamper spatial expansion. Finally to check if initially starting with templates in fewer number of sites can lead to the evolution and expansion of the protocell population even when protocells can degrade in the wet phase, we varied both the fraction of non-empty sites and the number of initial templates in them. We found that expansion is possible when there are at least 10% non-empty sites with at least 5 random templates in each of them and such expansion is possible even in presence of the protocell relocation process.

**Figure 6:**
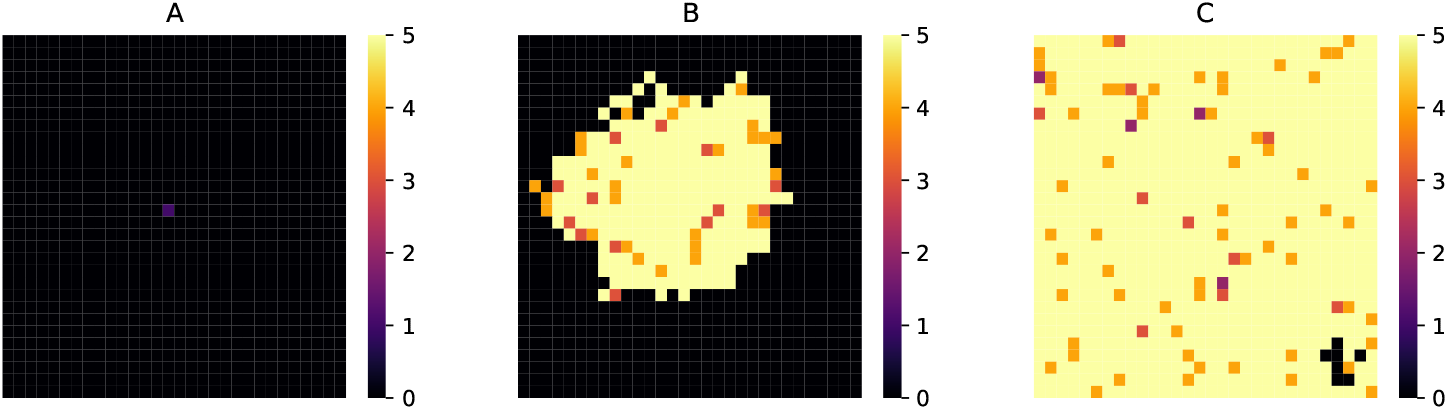
Expansion of protocell population from only one site containing 5 circular ssRNA templates initially. **A:** Initial stage; **B:** intermediate stage and **C:** when the percentage of sites with all 4 ribozymes becomes 90. Color values: Black:0 → no strands, Purple:1 → contains only non-enzymatic strands, Magenta:2 → contains 1 type of ribozyme, Orange:3 → contains 2 types of ribozymes, Dark Yellow:4 → contains 3 types of ribozymes, Light Yellow:5 → contains 4 types of ribozymes.

## 4 Discussion

Any compelling model of protocell evolution in a primordial RNA world must account for plausible environmental conditions in which such processes occurred and address the constraints imposed on the evolutionary processes by such conditions. The hot spring hypothesis of the origin of life provides an interesting test-bed to study evolutionary processes and assess the viability of such processes in aiding the formation, growth, division and eventual proliferation of protocells of increasing functional complexity. The ecological niche around hot-springs when subject to dry-wet cycling provides suitable conditions for the formation of long RNA polymers which can then be encapsulated into vesicles. Encapsulation of RNA sequences by protocells and replication of RNA via the rolling circle mechanism can lead to rapid growth of number of RNA strands and also increase the likelihood of creation of ribozymes of diverse functionality [32]. In this paper we examined how the coupling between the hot-spring environment and protocell evolution affects the proliferation of protocells containing multiple ribozymes. We showed how the interplay between the dry and the wet phase, where the former helps in polymerization reactions and the latter helps in protocell formation and division, can be critical for spreading the innovation in the form of ribozymes that are produced initially in a few protocells. In absence of dry-wet cycling such innovations are localized in a few regions and are likely to be lost in the course of evolution.

The hot spring hypothesis emphasizes the importance of the gel phase when protocells compartments can exchange molecules with each other by the strand diffusion process through their fused membranes. A question that can be raised in the light of our analysis is whether a gel phase characterized by diffusion of long RNA sequences is essential for the spatial proliferation of functionally complex protocells containing multiple ribozymes. It has been argued [35] that the usefulness of the gel phase stems from its ability to facilitate communal evolution. The fusion of the protocellular membranes lead to the creation of a single connected, albeit dispersed, community of RNA strands that can benefit from innovation in RNA sequences (such as appearance of a new ribozyme) that may appear in one region of the community. However, for such innovations to be effective, they need to be replicated and spread across the regions. Moreover, the advantage of communal evolution is considerably diminished if the communal structure is periodically destroyed due to environmental cycling. Our results indicate that diffusion of long RNA strands across fused protocell compartments in the gel phase inhibits the proliferation of functionally diverse protocells with the effect being most detrimental when the process of protocell degradation is taken into account.

Experimental studies [48, 51, 35] have already revealed the formation of multi-lamellar structures in the gel phase and budding of lipid vesicles containing DNA. The effectiveness of the rolling circle replication process [28, 32] used in our model to generate new RNA strands, especially in the presence of a polymerase ribozyme has recently been demonstrated [62]. Certain aspects of the hot-spring hypothesis captured in our computational model, such as the inhibitory effect of long RNA strand diffusion in the gel phase can be subjected to experimental validation in the lab. We therefore hope our work will stimulate further experimental investigations in the lab by simulating conditions prevalent near geothermal hot-springs and subject our model of protocell evolution to rigorous testing. The puzzle of origin of life is still far from being solved, but many pieces of the puzzle are beginning to fall into place. We believe that conceptual advances accompanied by new insights from experiments will enable us to make significant progress in understanding this challenging topic.

## Notes

### Competing Interest Statement

The authors have declared no competing interest.

