## Supplementary information for "The effect of environment on the evolution and proliferation of protocells of increasing complexity"

July 14, 2022

### 1 Supplementary videos

#### 1.1 Vid_S1

This video shows the evolution of the protocell population across the entire region for the case when gel phase diffusion is absent and protocell degradation does not occur. Heat maps for the entire lattice, denoting protocells with varying number of ribozymes, are generated at the beginning of each dry phase, after each dry phase and after each wet phase. Color coding: *black* → 0 → no strands, *purple* → 1 → contains only non-enzymatic strands, *magenta* → 2 → contains 1 type of ribozyme, *orange* → 3 → contains 2 types of ribozymes, *dark yellow* → 4 → contains 3 types of ribozymes, *light yellow* → 5 → contains 4 types of ribozymes.

#### 1.2 Vid_S2

This video shows the evolution of the protocell population across the entire region for the case when gel phase diffusion is present ($D = 1.28\ \mu m^2 h^{-1}$) and protocell degradation does not occur in the wet phase. Heat maps for the entire lattice, denoting protocells with varying number of ribozymes, are generated at the beginning of each dry phase, beginning of each wet phase and beginning of each gel phase. Color coding: *black* → 0 → no strands, *purple* → 1 → contains only non-enzymatic strands, *magenta* → 2 → contains 1 type of ribozyme, *orange* → 3 → contains 2 types of ribozymes, *dark yellow* → 4 → contains 3 types of ribozymes, *light yellow* → 5 → contains 4 types of ribozymes.

#### 1.3 Vid_S3

This video shows the evolution of the protocell population across the entire region for the case when wet phase protocell relocation is implemented by randomly reshuffling the position of each protocell at the beginning of each wet phase. We also assume that diffusion in the gel phase is absent and protocell degradation does not occur in the wet phase. Heat maps are generated at the beginning of each dry phase, beginning of each wet phase and after protocell relocation happens (but before protocell division). Color coding: *black* → 0 → no strands, *purple* → 1 → contains only non-enzymatic strands, *magenta* → 2 → contains 1 type of ribozyme, *orange* → 3 → contains 2 types of ribozymes, *dark yellow* → 4 → contains 3 types of ribozymes, *light yellow* → 5 → contains 4 types of ribozymes.

#### 1.4 Vid_S4

This video shows the time evolution of the dividing protocells and protocells containing either 3 or 4 types of ribozymes for the case when relocation of protocells in the wet phase is not allowed, diffusion in the gel phase is absent and protocell degradation does not occur in the wet phase. Color coding: *black* → 0 → protocells that are not dividing and contains less than 3 types of ribozymes, *purple* → 1 → non-dividing protocells containing any 3 of the 4 types of ribozymes, *magenta* → 2 → non-dividing protocells containing 4 types of ribozymes, *orange* $\rightarrow 3 \rightarrow$ dividing protocells containing any 3 of the 4 types of ribozymes, *light yellow* $\rightarrow 4 \rightarrow$ dividing protocells containing 4 types of ribozymes. For convenience of watching, the video is truncated in a way so that it starts when there is at least one protocell with 3 types of ribozymes.

#### 1.5 Vid_S5

This video shows the time evolution of the locations of dividing protocells and protocells containing multiple types of ribozymes for the case when wet phase protocell relocation is implemented by randomly reshuffling the position of each protocell at the beginning of each wet phase. We also assume that diffusion in the gel phase is absent and protocell degradation does not occur in the wet phase. Color coding: *black* $\rightarrow 0 \rightarrow$ protocells that are not dividing contains less than 3 types of ribozymes, *purple* $\rightarrow 1 \rightarrow$ non-dividing protocells containing any 3 out of the 4 types of ribozymes, *magenta* $\rightarrow 2 \rightarrow$ non-dividing protocells containing 4 types of ribozymes, *orange* $\rightarrow 3 \rightarrow$ dividing protocells containing any 3 out of the 4 types of ribozymes, *light yellow* $\rightarrow 4 \rightarrow$ dividing protocells containing 4 types of ribozymes. For convenience of watching, the video is truncated in a way so that it starts when there is at least one protocell with 3 types of ribozymes.

#### 1.6 Vid_S6

This video shows the evolution of the protocell population across the entire region for the case when gel phase diffusion is absent and protocell degradation occurs in the wet phase with a probability $P_{kill} = 0.01$. Heat maps are generated at the beginning of each dry phase, after each dry phase and after each wet phase. Color coding: *black* $\rightarrow 0 \rightarrow$ no strands, *purple* $\rightarrow 1 \rightarrow$ contains only non-enzymatic strands, *magenta* $\rightarrow 2 \rightarrow$ contains 1 type of ribozyme, *orange* $\rightarrow 3 \rightarrow$ contains 2 types of ribozymes, *dark yellow* $\rightarrow 4 \rightarrow$ contains 3 types of ribozymes, *light yellow* $\rightarrow 5 \rightarrow$ contains 4 types of ribozymes.

#### 1.7 Vid_S7

This video shows the evolution of the protocell population across the entire region for the case when diffusion of long RNA strands are allowed to occur in the gel phase ($D = 1.28\ \mu m^2 h^{-1}$) and protocell degradation occurs in the wet phase with probability $P_{kill} = 0.01$. Heat maps are generated at the beginning of each dry phase, beginning of each wet phase and beginning of each gel phase. Color coding: *black* $\rightarrow 0 \rightarrow$ no strands, *purple* $\rightarrow 1 \rightarrow$ contains only non-enzymatic strands, *magenta* $\rightarrow 2 \rightarrow$ contains 1 type of ribozyme, *orange* $\rightarrow 3 \rightarrow$ contains 2 types of ribozymes, *dark yellow* $\rightarrow 4 \rightarrow$ contains 3 types of ribozymes, *light yellow* $\rightarrow 5 \rightarrow$ contains 4 types of ribozymes.

### 2 Pseudo-code

**Note:** Periodic boundary condition are imposed after each time step and vesicle division event

**Initial setup:**

Lattice size $30 \times 30 = 900$ (N=30)
$K_{fast} = 0.362\ h^{-1}$
$K_{cyc} = 0.362\ h^{-1}$
Degradation rate $h = 0.0008\ hr^{-1}$
$P_r, P_c, P_n, P_p = 0.03, 0.03, 0.03, 0.03$
Create a NxN matrix for threshold volumes $V_T$ where each element is 100
Create a NxN matrix for S where each element is $0.8 \times 100$
Create a NxN matrix for $S_{max}$ where each element is $0.8 \times 100$
Create a matrix named cells, under which there are NxN empty lists denoting each cell
Fill each cell with 7 empty lists which will contain 7 types RNA strands described in our model
Fill the list for circ ssRNA of each cell with rates randomly assigned as,

$$K_{rep} \sim e^{-3.22-0.005(L-8)-2.8\times rand.norm(0.35,0.0667)}\ h^{-1}$$

Create an empty matrix 'tot-rate' which will contain the total rates $K^i_{tot}$ for each cell at any time step

**Time evolution:**

Begin loop over day

- **Dry phase:**
- Count the total number of nucleotide synthase ($n_o$) across all sites

  **Determine time step size:**

- Start loop over lattice sites
  - Calculate rate reduction factor $f_{ij} = (S_{ij} + bn_{ij})/S^{ij}_{max}$
  - If total number of strands in this site < 1000, create a list containing the rates for 6 types of reactions:

$$a = \left[ \sum_{k}^{s_{ij}} K^k_{rep} f_{ij}, \sum_{k}^{d_{ij}} K^k_{rep} f_{ij}, K_{fast} \frac{r_{ij} s_{ij} f_{ij}}{100}, K_{fast} \frac{r_{ij} d_{ij} f_{ij}}{100}, K_{cyc} \frac{c_{ij} l_{ij}}{100}, h(s_{ij}+d_{ij}+l_{ij}+r_{ij}+c_{ij}+n_{ij}+p_{ij}) \right]$$

  - Else create a list:

$$a = \left[ \sum_{k}^{s_{ij}} K^k_{rep} f_{ij} \, , \, 0 \, , \, K_{fast} \frac{r_{ij} s_{ij} f_{ij}}{100} \, , \, 0 \, , \, K_{cyc} \frac{c_{ij} l_{ij}}{100} \, , \, h(s_{ij} + d_{ij} + l_{ij} + r_{ij} + c_{ij} + n_{ij} + p_{ij}) \right]$$

  - Assign $tot - rate_{ij} = sum(a)$
- End loop over sites
- Find $K^{max}_{tot} = \max(\text{tot-rate})$
- Calculate $dt = 1/K^{max}_{tot}$
- Change time t=t+dt until $t \leq T_{dry}$

  **Reactions:**

- Create a counter variable (g) to count total number of replications
- Start loop over cells
  - If $K^{ij}_{tot} > 0$ and $random.uniform(0, 1) < K^{ij}_{tot} \times dt$
    * Calculate rate reduction factor $f_{ij} = (S_{ij} + bn_{ij})/S^{ij}_{max}$
    * If total number of strands in this site < 1000, create a list containing the rates for 6 types of reactions:

$$a = \left[ \sum_{k}^{s_{ij}} K^k_{rep} f_{ij} \, , \, \sum_{k}^{d_{ij}} K^k_{rep} f_{ij} \, , \, K_{fast} \frac{r_{ij} s_{ij} f_{ij}}{100} \, , \, K_{fast} \frac{r_{ij} d_{ij} f_{ij}}{100} \, , \, K_{cyc} \frac{c_{ij} l_{ij}}{100} \, , \right.$$
$$\left. h(s_{ij} + d_{ij} + l_{ij} + r_{ij} + c_{ij} + n_{ij} + p_{ij}) \right]$$

    * Else create a list:

$$a = \left[ \sum_{k}^{s_{ij}} K^k_{rep} f_{ij} \, , \, 0 \, , \, K_{fast} \frac{r_{ij} s_{ij} f_{ij}}{100} \, , \, 0 \, , \, K_{cyc} \frac{c_{ij} l_{ij}}{100} \, , \right.$$
$$\left. h(s_{ij} + d_{ij} + l_{ij} + r_{ij} + c_{ij} + n_{ij} + p_{ij}) \right]$$

    * Choose the type of reaction randomly based the relative propensities $(a/K^i_{tot})$ of the reactions
    * If reaction is circ ssRNA to circ dsRNA

- Choose a circ ssRNA randomly from the list of circ ssRNA based their relative $K_{rep}$ values
- Add a $K_{rep}$ to the list of circ dsRNA by multiplying the chosen $K_{rep}$ value with $e^{-0.005L+0.005(L-8)}$
- Remove the chosen $K_{rep}$ from the list of circ ssRNA
- Add +1 to g

* If reaction is circ dsRNA to open-ended ssRNA
  - Draw a random number $p = random.uniform(0, 1)$
  - If $p < P_r$ then add a 0 to the list of replicase
  - If $P_r \geq p < P_r + P_c$ then add a 0 to the list of cyclase
  - If $P_r + P_c \geq p < P_r + P_c + P_n$ then add a 0 to the list of nucleotide synthase
  - If $P_r + P_c + P_n \geq p < P_r + P_c + P_n + P_p$ then add a 0 to the list of peptidyl transferase
  - If $p \geq P_r + P_c + P_n + P_p$ then add a 0 to the list of open-ended ssRNA
  - add +1 to g
* If reaction is replicase catalyzed circ ssRNA to circ dsRNA
  - Choose a circ ssRNA randomly from the list of circ ssRNA
  - Add a $K_{rep}$ to the list of circ dsRNA by multiplying the chosen $K_{rep}$ value with $e^{-0.005L+0.005(L-8)}$
  - Remove the chosen $K_{rep}$ from the list of circ ssRNA
  - Add +1 to g
* If reaction is replicase catalyzed circ dsRNA to open-ended ssRNA
  - Draw a random number $p = random.uniform(0, 1)$
  - If $p < P_r$ then add a 0 to the list of replicase
  - If $P_r \geq p < P_r + P_c$ then add a 0 to the list of cyclase
  - If $P_r + P_c \geq p < P_r + P_c + P_n$ then add a 0 to the list of nucleotide synthase
  - If $P_r + P_c + P_n \geq p < P_r + P_c + P_n + P_p$ then add a 0 to the list of peptidyl transferase
  - If $p \geq P_r + P_c + P_n + P_p$ then add a 0 to the list of open-ended ssRNA
  - add +1 to g
* IF reaction is cyclase catalyzed open-ended ssRNA to circ ssRNA
  - Add a random $K_{rep}$ value to the list of circ ssRNA following:

$$K_{rep} \sim e^{-3.22-0.005(L-8)-2.8\times rand.norm(0.35,0.0667)}\ h^{-1}$$

  - Remove a $K_{rep}$ value from the list of circ ssRNA
* If reaction is degradation
  - Choose which type of RNA strand will lose one member, randomly with probabilities proportional to the number of each type in this cell
  - Go the list of the chosen type and remove one member randomly from that list
* After reaction if $p_{ij} > 0$
  - Change $V_T^{ij}$ as $V_T^{ij} = 100 + 20p_{ij}$

- End loop over sites
- Calculate $S_{tot} = \sum_{ij} S_{ij} + (bn_o - g)$
- Start loop over sites
  - Assign $S_{ij} = (S_{tot}/N^2)$
  - If new $S_{ij} < 0$, put $S_{ij} = 0$
  - If new $S_{ij} > S_{max}^{ij}$, put $S_{ij} = S_{max}^{ij}$
- End loop over sites

**Wet phase:**

- Randomly shuffle the protocell positions on the lattice by using Fisher-Yates algorithm
- Start loop over protocells
  - If random.uniform(0,1) $< P_{kill}$ then make this protocell empty
- End loop over protocells
- Create a list of protocell indices
- Randomly shuffle the indices
- Start loop over indices
  - Count total number of strands $(s_{ij} + d_{ij} + l_{ij} + r_{ij} + c_{ij} + n_{ij} + p_{ij})$ inside the cell
  - If $(s_{ij} + d_{ij} + l_{ij} + r_{ij} + c_{ij} + n_{ij} + p_{ij}) \geq V_T^{ij}$
    * Create an empty list for weakness
    * Start loop over 8 nearest neighbors
      · If cell index is not marked 'X' and total number of strands inside it $(s_{i'j'}+d_{i'j'}+l_{i'j'}+r_{i'j'}+c_{i'j'}+n_{i'j'}+p_{i'j'})$ is lower than that of the dividing cell, then Assign weakness $w_{i'j'} = (s_{ij}+d_{ij}+l_{ij}+r_{ij}+c_{ij}+n_{ij}+p_{ij})-(s_{i'j'}+d_{i'j'}+l_{i'j'}+r_{i'j'}+c_{i'j'}+n_{i'j'}+p_{i'j'})$

      · Else assign weakness $w_{i'j'} = 0$
    * Choose a cell randomly with probabilities equal to the relative weaknesses of the cells
    * Make the chosen cell empty
    * Mark the cell index for both the dividing cell and the eliminated cell 'X'
    * Start loop over each strand of the dividing cell
      · Draw a random number p

      · If $p < 0.5$ move the strand to the empty cell and remove from the present cell

      · Else keep the strand in the present cell
    * End loop over strands
    * Change the $V_T$ of the daughter cells as $V_T^{ij} = 100 + 20p_{ij}$ and $V_T^{i'j'} = 100 + 20p_{i'j'}$
- End loop over indices

**Gel phase:**

- Start loop over diffusion time steps
  - Create a list of site indices
  - Randomly shuffle the indices
  - Start loop over indices
    * Draw a molecule randomly from this site M times, where M is total number of strands in this site
    * After each draw choose a neighbor randomly from 8 nearest neighbors
    * If the neighbor has fewer number of strands and random.uniform(0,1) $< P_{hop}$ then move that strand to its corresponding list in the neighbor
  - End loop over indices
- End loop over diffusion time steps

End loop over days
